# GenomegaMap: within-species genome-wide *d*_*N*_/*d*_*S*_ estimation from over 10,000 genomes

**DOI:** 10.1101/523316

**Authors:** Daniel J. Wilson, The CRyPTIC Consortium

**Author notes:** Address for correspondence: Big Data Institute, Nuffield Department of Population Health, Li Ka Shing Centre for Health Information and Discovery, Oxford, OX3 7LF, United Kingdom. ORCID: 0000-0002-0940-3311 (Daniel J. Wilson). For consortium member list, see Appendix A (The CRyPTIC Consortium).

## Abstract

The *d*_*N*_/*d*_*S*_ ratio provides evidence of adaptation or functional constraint in protein-coding genes by quantifying the relative excess or deficit of amino acid-replacing versus silent nucleotide variation. Inexpensive sequencing promises a better understanding of parameters such as *d*_*N*_/*d*_*S*_, but analysing very large datasets poses a major statistical challenge. Here I introduce genomegaMap for estimating within-species genome-wide variation in *d*_*N*_/*d*_*S*_, and I apply it to 3,979 genes across 10,209 tuberculosis genomes to characterize the selection pressures shaping this global pathogen. GenomegaMap is a phylogeny-free method that addresses two major problems with existing approaches: (i) it is fast no matter how large the sample size and (ii) it is robust to recombination, which causes phylogenetic methods to report artefactual signals of adaptation. GenomegaMap uses population genetics theory to approximate the distribution of allele frequencies under general, parent-dependent mutation models. Coalescent simulations show that substitution parameters are well-estimated even when genomegaMap’s simplifying assumption of independence among sites is violated. I demonstrate the ability of genomegaMap to detect genuine signatures of selection at antimicrobial resistance-conferring substitutions in *M. tuberculosis* and describe a novel signature of selection in the cold-shock DEAD-box protein A gene *deaD/csdA*. The genomegaMap approach helps accelerate the exploitation of big data for gaining new insights into evolution within species.

---

Interpreting patterns of substitution in genetic sequences is a fundamental approach in evolutionary biology. For example, an excess rate of amino acid-replacing *non-synonymous* substitution compared to silent *synonymous* substitution, quantified by the *d*_*N*_/*d*_*S*_ ratio (also denoted *K*_*A*_/*K*_*S*_ or *ω*), provides evidence of adaptive evolution, while the reverse pattern, more prevalent in functional protein-coding sequences, provides evidence for purifying selection (e.g. Miyata and Yasunaga 1980; *Perler et al.* 1980; Nei and Gojobori 1986; Nielsen and Yang 1998).

However, estimating substitution parameters typically relies on first estimating, or co-estimating, a phylogenetic tree relating the observed sequences. Two major drawbacks commonly arise when (i) recombination is present or (ii) sample sizes are large. The first major drawback, often encountered in analyses of within-species variation, is that recombination breaks the assumption of a single phylogeny, and instead generates a network of ancestral relationships in which different genes, and different positions within genes, can have different phylogenetic histories (Schierup and Hein 2000). It is well established that inappropriate application of phylogeny-based methods to recombining data can produce highly misleading biological inferences, including false signals of adaptive evolution in the form of artificially elevated *d*_*N*_/*d*_*S*_ (Anisimova *et al.* 2003; Shriner *et al.* 2003).

The second major drawback is the computational cost of estimating a phylogeny when the number of sequences becomes large, for example the 10,209 genomes recently published by The CRyPTIC Consortium and The 100,000 Genomes Project (2018) that bear witness to the relentless evolution of antimicrobial resistance in tuberculosis. This is a double blow because the cost of evaluating the fit of an individual phylogeny increases at the same time as the number of possible phylogenies explodes (Felsenstein 1973, 1978). The problem will become increasingly acute with the steady march towards ever more sequencing.

Wilson and McVean (2006) developed a method, *omegaMap*, to estimate *d*_*N*_/*d*_*S*_ in the presence of recombination. While *omegaMap* avoids the false signals of adaptive evolution suffered by phylogenetic methods, its application to large datasets is limited by the underlying PAC (product of approximate conditionals) approach, whose computational complexity increases quadratically with sample size (Li and Stephens 2003).

In this paper I address these drawbacks with existing methods by introducing *genomegaMap*, a phylogeny-free statistical approach to estimating substitution parameters that implicitly integrates over phylogenetic relatedness using diffusion theory and the coalescent (Wright 1949; Kingman 1982). The computational cost of the method remains constant even as the sample size increases arbitrarily, making it a viable approach for extremely large datasets. The method assumes independence between sites, yet simulations show that the method performs well even when the absence of recombination causes strong linkage disequilibrium. I demonstrate the utility of the method by estimating variation in *d*_*N*_/*d*_*S*_ ratios in 3,979 genes sequenced in 10,209 *M. tuberculosis* genomes (The CRyPTIC Consortium and The 100,000 Genomes Project 2018).

## Methods

### Population Genetics Model

Estimating the *d*_*N*_/*d*_*S*_ ratio can be seen as a special case of the more general problem of estimating a substitution rate matrix. The Nielsen and Yang (1998) (NY98) codon model assumes that a non-synonymous substitution occurs at *ω* times the rate of its synonymous counterpart. It is defined by the following substitution rate from codon *i* to *j* (*j* ≠ *i*):

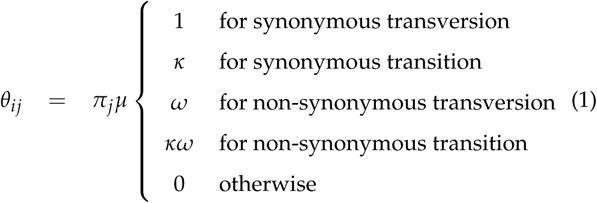

where *ω* is the *d*_*N*_/*d*_*S*_ ratio, *κ* the transition:transversion ratio and *π*_*j*_ the equilibrium frequency of allele *j*. To form a proper rate matrix, the diagonal elements must be defined as *θ*_*ii*_ = − Σ_*j*≠*i*_ *θ*_*ij*_. The scaling constant *μ* is determined by the expected substitution rate, *θ* = − Σ_*i*_ Σ_*j*≠*i*_ *π*_*i*_ *θ*_*ij*_. Following the convention in population genetics, the rate is defined in units of 2*PN*_*e*_ generations, where *P* is the ploidy and *N*_*e*_ the effective population size.

*GenomegaMap* estimates substitution parameters by modeling the allele frequency distribution at each site. Analyses of *d*_*N*_/*d*_*S*_ within species (e.g. Nielsen and Yang 1998; Wilson and McVean 2006) have implicitly treated selection as a form of *mutational bias*, in which the mutation rate matrix equals the NY98 substitution rate matrix, and fitness differences between individuals are ignored. I follow the convention here. (For an alternative approach, see *gammaMap* (Wilson et al. 2011), which separately models mutation and selection.)

The distribution of allele frequencies under the simplifying assumptions of a stable and unstructured population, selective neutrality, and *parent independent* mutation, in which the rate of mutation from allele *i* to *j*, *θ*_*ij*_ = *θ*_*·j*_ depends only on the target allele, *j*, is derived from diffusion theory and attributed to Wright (1949) (see, e.g.,Watterson 1977):

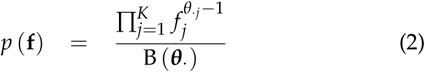

where *f*_*j*_ is the frequency of allele *j*, *K* is the number of alleles and 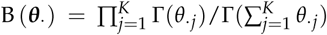 is the multivariate beta function.

For more general, *parent-dependent*, mutation models, the distribution cannot be easily calculated. Instead, I employ the approach of Wilson *et al.* (2011, Equation B1) who approximated the conditional allele frequency distribution given the identity of the oldest allele *A* as a Dirichlet distribution, so that

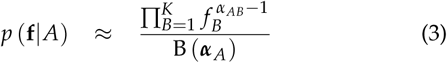

where *α*_*AB*_ = *m*_*AB*_/*m*_*AA*_ and *m*_*AB*_ is the probability of sampling an allele *B* conditional on having sampled allele *A* in a sample of size two, calculable using the coalescent as

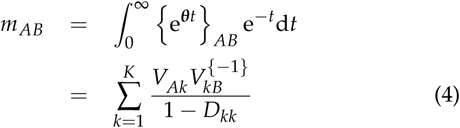

where ***θ*** = ***VDV***^−1^ is the eigen decomposition of the substitution rate matrix. This approximation, which in principle allows any Markovian substitution process to be fitted, is motivated by a low mutation rate assumption and therefore expected to work best when the expected number of substitutions per site is small.

Assuming random sampling, the conditional allele count distribution is Multinomial-Dirichlet distributed, so that

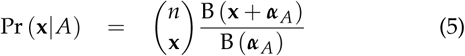

where *x*_*j*_ is the number of times allele *j* was counted and *n* the sample size. The identity of the oldest allele *A* is then averaged over to obtain a likelihood for the allele count:

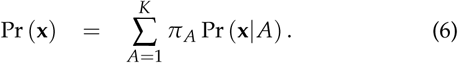

The coarsest approximation made by *genomegaMap* is independence between sites, which is motivated by the benefits it confers with the rest of the model: (i) The computational complexity is constant irrespective of sample size, whereas the likelihoods in phylogenetic and PAC models increase linearly and quadratically with sample size, respectively. (ii) Missing data can be handled easily because the sample size need not be the same from site-to-site. (iii) No haplotype information is required.

### Simulations

I performed simulations to test the performance of *genomegaMap* under two scenarios. In the Unlinked simulations, every codon was simulated independently, in keeping with the assumption of *genomegaMap*. In the Clonal simulations, all codons were completely linked, maximally violating this assumption of *genomegaMap*. For each scenario, I simulated 100 datasets of 334 codons in 10,000 individuals. The parameters were simulated independently for each dataset from log-normal distributions with (2.5%, 97.5%) quantiles of (0.05, 5) for *ω*, (1, 9) for *κ* and (0.001, 0.1) for *θ*. *ω* was assumed constant along the sequence. Codon frequencies were simulated from the empirical codon frequency distribution among 10,209 *M. tuberculosis* genomes (The CRyPTIC Consortium and The 100,000 Genomes Project 2018). For each simulated dataset, parameters were estimated by Markov chain Monte Carlo (MCMC), using as priors the same distributions used to simulate *ω*, *κ* and *θ*. Under these conditions, the 95% credibility intervals (CIs) should include the true parameters in 95% of simulations, if the approximate likelihood performs optimally (Dawid 1982). For each analysis I ran two independent MCMC chains of 10,000 iterations.

### *Analysis of* Neisseria meningitidis porB3

To compare *genomegaMap* to *omegaMap*, I reanalysed 23 of 79 *porB3 N. meningitidis* sequences of Urwin *et al.* (2002) comprising the *carriage study* subset of Wilson and McVean (2006). Columns in the alignment with any indels were removed to aid the comparison because *omegaMap* handles them differently. I assumed an exponential prior distribution with mean 1.0 for *ω* and improper log-uniform priors for *κ* and *θ*. I assumed a Bayesian sliding window (i.e. piecewise constant) model for variation in *ω* along the gene, with a mean window length of 30 codons (Wilson and McVean 2006). For both *genomegaMap* and *omegaMap*, I ran two independent MCMC chains of 500,000 iterations.

### Analysis of 10,209 Mycobacterium tuberculosis Genomes

The CRyPTIC Consortium and The 100,000 Genomes Project (2018) collected and whole-genome sequenced 10,209 *M. tuberculosis* samples from 16 countries across six continents comprising strains enriched for antimicrobial resistance and unenriched strains collected for routine clinical diagnostics. They mapped all genomes to the H37Rv reference genome (Cole *et al.* 1998) (Genbank accession number NC_000962.2). I downloaded the alignment of every genome to H37Rv and combined these to create a multiple sequence alignment for each of the 3,979 coding sequences (CDSs) in the Genbank annotation, ignoring insertions relative to H37Rv and masking nonsense mutations.

Inference of *ω*, *κ* and *θ* for an individual gene can be improved by gleaning information from other genes. Often this is implemented through a hierarchical model, for example estimating a distribution for the selection parameters across all sites in all genes (Wilson *et al.* 2011). However, hierarchical modeling requires sophisticated techniques for simultaneously analysing thousands of genes across a high performance computing cluster. Instead, I mimicked a hierarchical model heuristically by training a prior for *ω*, *κ* and *θ* using an alignment of 334 codons randomly chosen from the 3,979 genes. For this preliminary analysis, I employed an exponential hyperprior with mean 1.0 for *ω*, imposing a single window across the alignment, and improper log-uniform hyperpriors for *κ* and *θ*, running two MCMC chains for 10,000 iterations. This produced posterior means of −0.79, 1.2 and −2.9 and standard deviations of 0.20, 0.21 and 0.15 for log *ω*, log *κ* and log *θ* respectively.

I used these results to form priors for the analyses of the 3,979 individual genes by assuming log-normal distributions, multiplying the standard deviation parameters by 10 for *ω* and 3.2 for *κ* and *θ* to avoid over-informative priors. This produced a prior median and (2.5%, 97.5%) quantiles of 0.45 (0.0098, 21) for *ω*, 3.2 (0.90, 12) for *κ* and 0.057 (0.023, 0.14) for *θ*. I used the genome-wide empirical codon frequency distribution and assumed a Bayesian sliding window model for variation in *ω* along each gene, with a mean window length of 33 codons. For each gene I ran two independent MCMC chains of 500,000 iterations.

### Software and Data Availability

*GenomegaMap* is available as a Docker container and C++ source code from https://hub.docker.com/r/dannywilson/gcat-omegamap and https://github.com/danny-wilson/gcat-omegaMap. The following data are available: codon counts for every annotated coding sequence https://doi.org/10.6084/m9.figshare.7599020.v1, figures illustrating variation in *ω* along every coding sequence https://doi.org/10.6084/m9.figshare.7599029.v1, and summaries of the posterior distribution of *ω*, *κ* and *θ* for every coding sequence https://doi.org/10.6084/m9.figshare.7599032.v1.

## Results and Discussion

### *General Performance of* GenomegaMap

The motivation for developing *genomegaMap* came from the observation that *omegaMap* estimates of substitution parameters, including the *d*_*N*_/*d*_*S*_ ratio *ω*, were not strongly affected by the exact value of the recombination rate, as long as it was non-zero. This observation is reflected in the comparison of the analyses of the *N. meningitidis porB3* gene (Figure 1), for which the point estimates and 95% CIs of *ω* were almost identical between *omegaMap* and *genomegaMap*, even though the latter assumes codons are independent, i.e. unlinked. While the results were near-identical, the *genomegaMap* point estimates and 95% CIs were slightly more conservative, in the sense that they were closer to the prior expectation of *ω* = 1. These results suggest that substitution parameters are well-estimated within species when sites are assumed independent, despite the presence of linkage disequilibrium.

**Figure 1.**
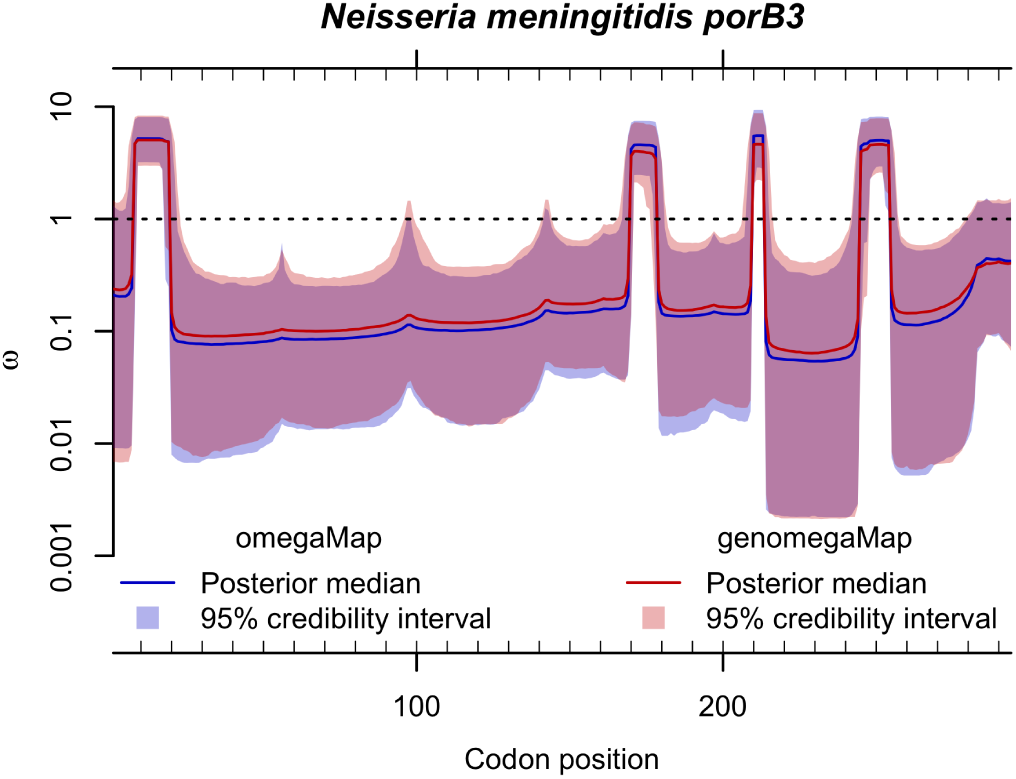
Comparison of *omegaMap* and *genomegaMap* estimates of the *d*_*N*_/*d*_*S*_ ratio *ω* along the *porB3* outer membrane protein gene of *Neisseria meningitidis*. Solid lines and shaded regions show the point estimates (posterior medians) and 95% credibility intervals respectively for *omegaMap* (in blue) and *genomegaMap* (in red). The *genomegaMap* analysis was 4.9 times faster for these 23 sequences.

To test this claim more thoroughly, I evaluated the relative performance of *genomegaMap* in two scenarios. In the Unlinked simulations, 334 codons were simulated independently across 10,000 individuals, favoring the *genomegaMap* assumption. In the Clonal simulations, all codons were completely linked, strongly violating the *genomegaMap* assumption of unlinked sites. As expected, *genomegaMap* performed well in the Unlinked simulations, producing point estimates strongly correlated with the true values of the *d*_*N*_/*d*_*S*_ ratio *ω*, the transition:transversion ratio *κ* and the mutation rate *θ*, and 95% CIs that included the truth in 98%, 98% and 97% respectively of the 100 simulations (Figure 2).

**Figure 2.**
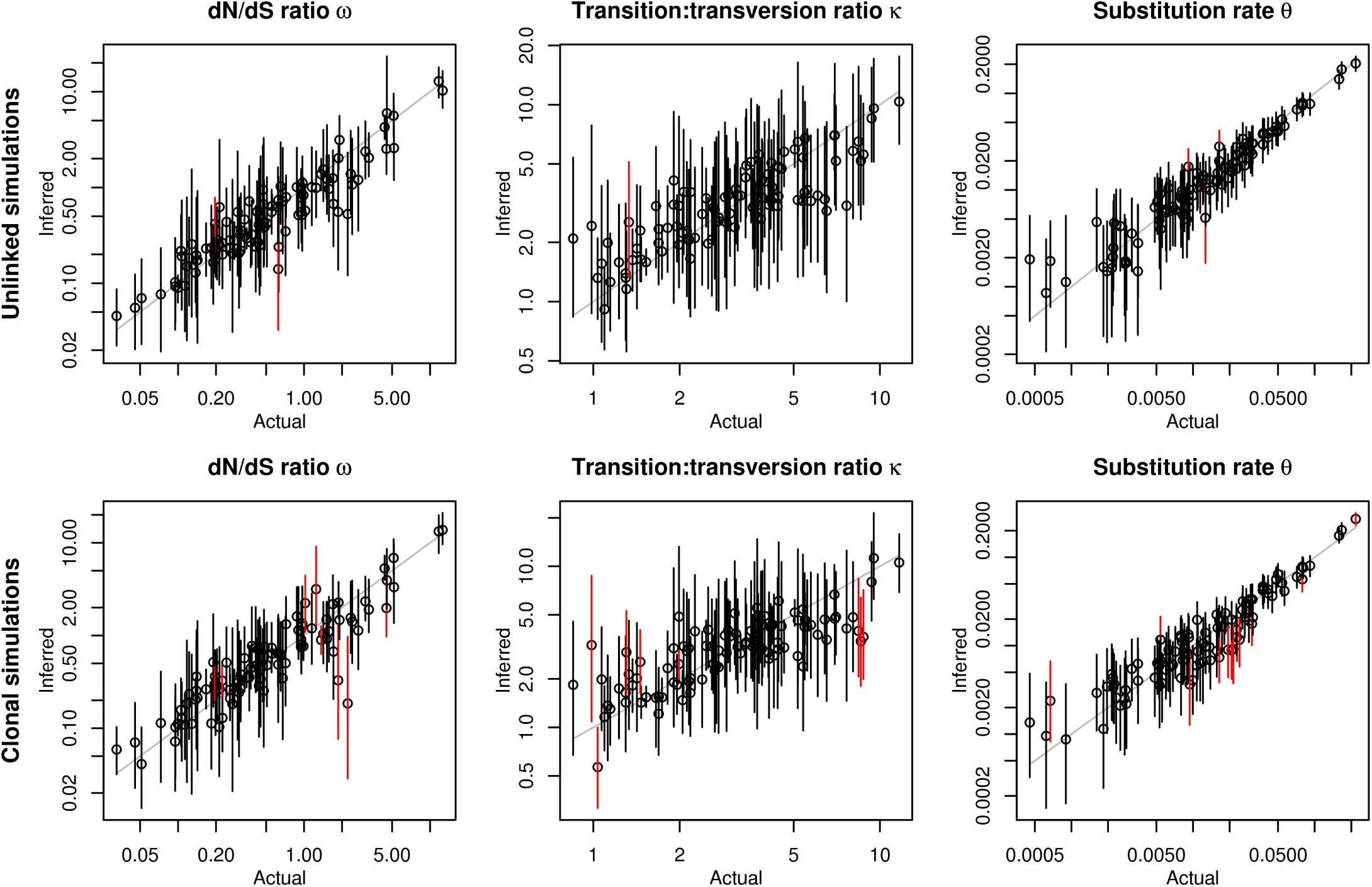
Performance of *genomegaMap* inference of *ω*, *κ* and *θ* in simulations. In the Unlinked simulations (top row) every codon was simulated independently, favoring the *genomegaMap* assumption. In the Clonal simulations (bottom row), all codons were completely linked, disfavoring the *genomegaMap* assumption. Point estimates (posterior medians) and 95% credibility intervals are indicated by the circles and solid vertical lines respectively, the latter colored red when they exclude the actual parameter. The number of simulations (out of 100) in which the 95% credibility intervals included the actual values of *ω*, *κ* and *θ* were 98, 98 and 97 in the Unlinked simulations and 92, 92 and 88 in the Clonal simulations. The correlation between the point estimates and actual values of log *ω*, log *κ* and log *θ* were 0.86, 0.69 and 0.92 in the Unlinked simulations and 0.82, 0.61 and 0.88 in the Clonal simulations.

In the Clonal simulations, codons were completely linked, maximally violating the independence assumption of *genomegaMap*. Despite this, the correlation between point estimates and true parameters remained strong, while the 95% CIs still included the truth in 92% of the 100 simulations for *ω* and *κ* and 88% of simulations for *θ* (Figure 2). These results suggest that *genomegaMap* produces only small loss in the accuracy of its point estimates and 95% CIs even when its independence assumption is completely wrong.

The major advantage of *genomegaMap* over *omegaMap* is its robustness to sample size. The computational run time of *omegaMap* increases with the square of the sample size. The run time of a comparable phylogenetic method would increase linearly with the sample size if the phylogeny were known; in practice co-estimating the phylogeny makes the computation much more intensive. In contrast, the run time of *genomegaMap* is constant with respect to sample size. This means it is uniquely suitable for the analysis of extremely large within-species data. To demonstrate its capabilities, I applied *genomegaMap* to 3,979 genes across 10,209 *M. tuberculosis* genomes.

### *Characterizing Selection in 10,209* M. tuberculosis *Genomes*

*Mycobacterium tuberculosis* is a bacterial pathogen responsible for tuberculosis, one of the world’s leading causes of death. 23% of the global population is thought to carry latent infection, of whom 9.0–11.1 million people are estimated to have developed tuberculosis in 2017, with 1.5–1.7 million resulting deaths. Drug resistance is a major problem for tuberculosis treatment; an estimated 483,000–639,000 new cases were resistant to first-line drugs in 2017 (World Health Organization 2018).

The aim of the CRyPTIC Consortium is to help improve control of tuberculosis and facilitate better, faster and more targeted treatment of drug-resistant tuberculosis via genetic resistance prediction, paving the way towards universal drug susceptibility testing. The CRyPTIC Consortium and The 100,000 Genomes Project (2018) collected and whole-genome sequenced 10,209 *M. tuberculosis* genomes to quantify the performance of genomic prediction of drug resistance. The predictions were correct in 91.3–97.5% of resistant isolates and 93.6–99.0% of susceptible isolates for the four first-line drugs.

These predictions rely on existing knowledge of the genetic mechanisms of drug resistance. Vast datasets have the potential to reveal novel mechanisms of drug resistance through genomewide association studies (GWAS). Such studies can benefit from an understanding of the selection pressures shaping genetic diversity and the identification of sites under positive selection because often that selection is driven by drug therapy (e.g. Pepperell *et al.* 2013; Zhang *et al.* 2013; Farhat *et al.* 2013; *Osório et al.* 2013; Lee *et al.* 2015; Koch *et al.* 2017; Mortimer *et al.* 2018).

*M. tuberculosis* is known for its complete lack, or near-complete lack, of homologous recombination (*Godfroid et al. 2018*), but as simulations showed, *genomegaMap* inference is robust to both recombination and the lack of recombination. I analysed the 3,979 genes sequenced across the 10,209 genomes with *genomegaMap*. Figure 3 summarizes the evidence for positive selection across the genome by quantifying the posterior probability of *ω >* 1. Most codons in most genes showed strong evidence against positive selection, i.e. Pr(*ω >* 1) ≪ 0.5, indicating strong functional constraint. Very few genes, such as *pncA* encoding pyrazinamidase, appeared to be dominated by positive selection. More often, the strongest evidence for positive selection was found in a very small number of codons within genes dominated by negative selection, such as *gyrA*, encoding DNA gyrase subunit A. This shows how positive selection occurs against backdrops of both rapid amino acid change and strong functional constraint, so the mean Pr(*ω >* 1) per gene provides limited insight.

**Figure 3.**
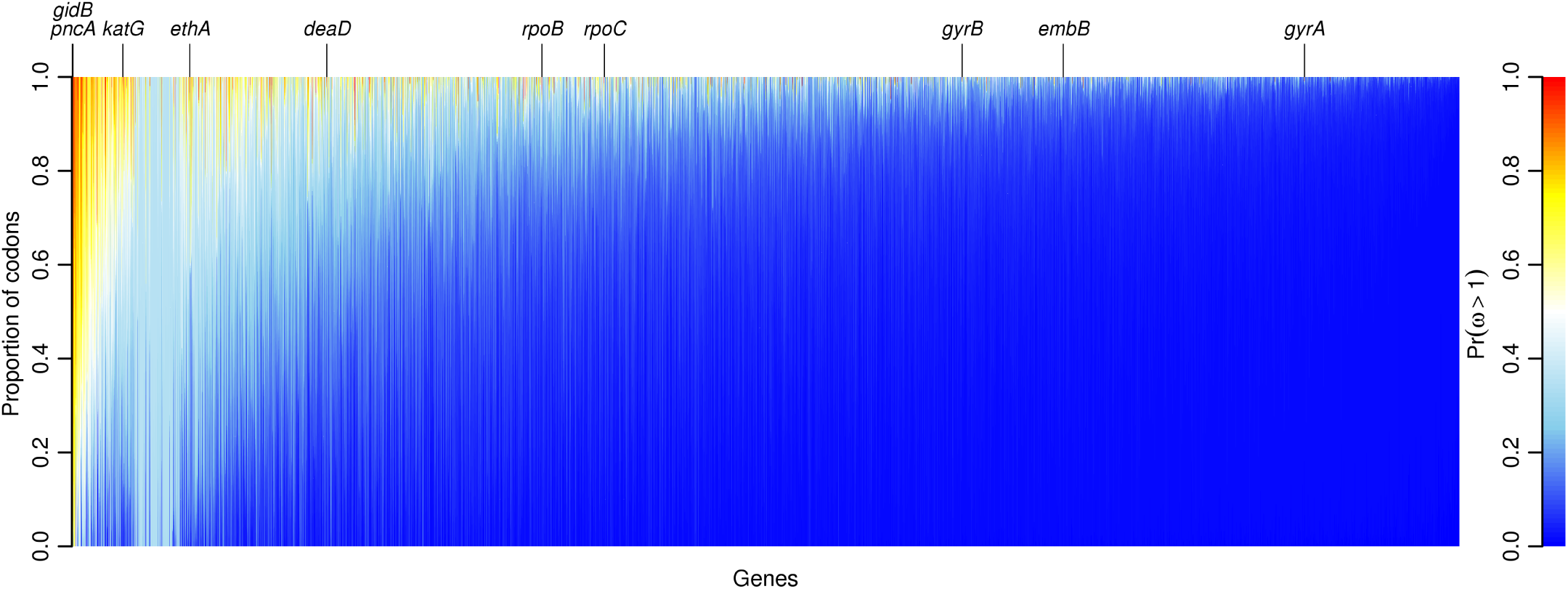
The evidence for positive selection across 3,979 genes in 10,209 *Mycobacterium tuberculosis* genomes. Each column is a stacked bar chart showing the proportion of codons in one gene with a given strength of evidence for positive selection, indicated by color. Blue indicates weakest evidence, Pr(*ω >* 1) ≠ 0, while red indicates strongest evidence, Pr(*ω >* 1) ≠ 1. Genes are ordered left-to-right by the mean Pr(*ω >* 1) across codons, from highest to lowest. Notable genes containing codons with strong evidence of positive selection are labeled; these occur throughout the spectrum. The block of genes with almost entirely sky blue coloration, roughly between *katG* and *ethA*, contained little information because they mapped poorly to the reference genome.

Instead, I identified every gene with one or more codons exhibiting a posterior probability of positive selection of at least 90% (i.e. Pr(*ω >* 1) ≥ 0.9), further classifying them by high, intermediate and low mean Pr(*ω >* 1) (Tables 1–3). The genes are annotated by their descriptions in GenBank or, when more informative, MycoBrowser (Kapopoulou *et al.* 2011). In total, 2,320/1,330,612 (0.2%) codons spanning 116/3,979 (3%) genes showed strong evidence of positive selection. Many occurred in genes encoding membrane proteins, toxin-antitoxin proteins (Sala *et al.* 2014), PE/PPE family proteins (Fishbein *et al.* 2015) and ESX family proteins (Gröschel *et al.* 2016).

**Table 1.** *Mycobacterium tuberculosis* genes with high mean Pr(*ω >* 1) and individual codons with Pr(*ω >* 1) *≥* 0.9

| Gene | Mean<br>$\Pr(\omega > 1)$ | Num. codons<br>$\Pr(\omega > 1) \geq 0.9$ | Product |
| --- | --- | --- | --- |
| gidB | 0.93 | 192 | 16S rRNA methyltransferase GidB |
| whiB6 | 0.92 | 81 | transcriptional regulatory protein WHIB-like WHIB6 |
| pncA | 0.91 | 135 | pyrazinamidase |
| Rv2621c | 0.83 | 6 | transcriptional regulator |
| furB | 0.79 | 46 | ferric uptake regulation protein FURB |
| Rv0456A | 0.79 | 26 | * possible toxin MazF1 |
| phoR | 0.71 | 289 | two component system response sensor kinase membrane associated |
| Rv1194c | 0.66 | 21 | HP <sup>a</sup> |
| Rv1672c | 0.65 | 135 | integral membrane transport protein |
| Rv0026 | 0.63 | 30 | HP |
| Rv2012 | 0.62 | 6 | HP |
| Rv3528c | 0.60 | 16 | HP |
| Rv0892 | 0.60 | 23 | monooxygenase |
| Rv1896c | 0.54 | 76 | HP |
| fgd2 | 0.51 | 4 | F420-dependent glucose-6-phosphate dehydrogenase |
| Rv0726c | 0.49 | 18 | * possible S-adenosylmethionine-dependent methyltransferase |
| Rv1830 | 0.47 | 69 | HP |
| Rv3294c | 0.43 | 11 | HP |
| pks6 | 0.43 | 239 | membrane bound polyketide synthase |
| Rv3847 | 0.43 | 5 | HP |
| Rv0229c | 0.40 | 5 | * possible conserved membrane protein with PIN domain |
| PE_PGRS15 | 0.38 | 7 | PE-PGRS family protein |
| Rv2100 | 0.37 | 7 | HP |
| Rv2706c | 0.37 | 4 | HP |
| Rv1129c | 0.37 | 10 | transcriptional regulator |
| cyp141 | 0.36 | 42 | cytochrome P450 141 |
| Rv2712c | 0.36 | 7 | HP |
| Rv2274c | 0.35 | 2 | * possible toxin MazF8 |
| Rv2787 | 0.34 | 23 | * conserved hypothetical alanine rich protein |
| Rv0246 | 0.34 | 31 | * probable conserved integral membrane protein |
| Rv0987 | 0.33 | 59 | adhesion component transport transmembrane protein |
| ethA | 0.33 | 17 | monooxygenase |
| fadD23 | 0.32 | 2 | acyl-CoA synthetase |
| Rv2035 | 0.32 | 7 | HP |
| Rv1507A | 0.31 | 11 | HP |
| Rv1265 | 0.31 | 33 | HP |
| Rv2438A | 0.31 | 3 | HP |
| pks8 | 0.30 | 13 | polyketide synthase |
| pks9 | 0.29 | 28 | polyketide synthase |

<sup>a</sup> HP: hypothetical protein. \* Mycobrowser annotation

**Table 2.**
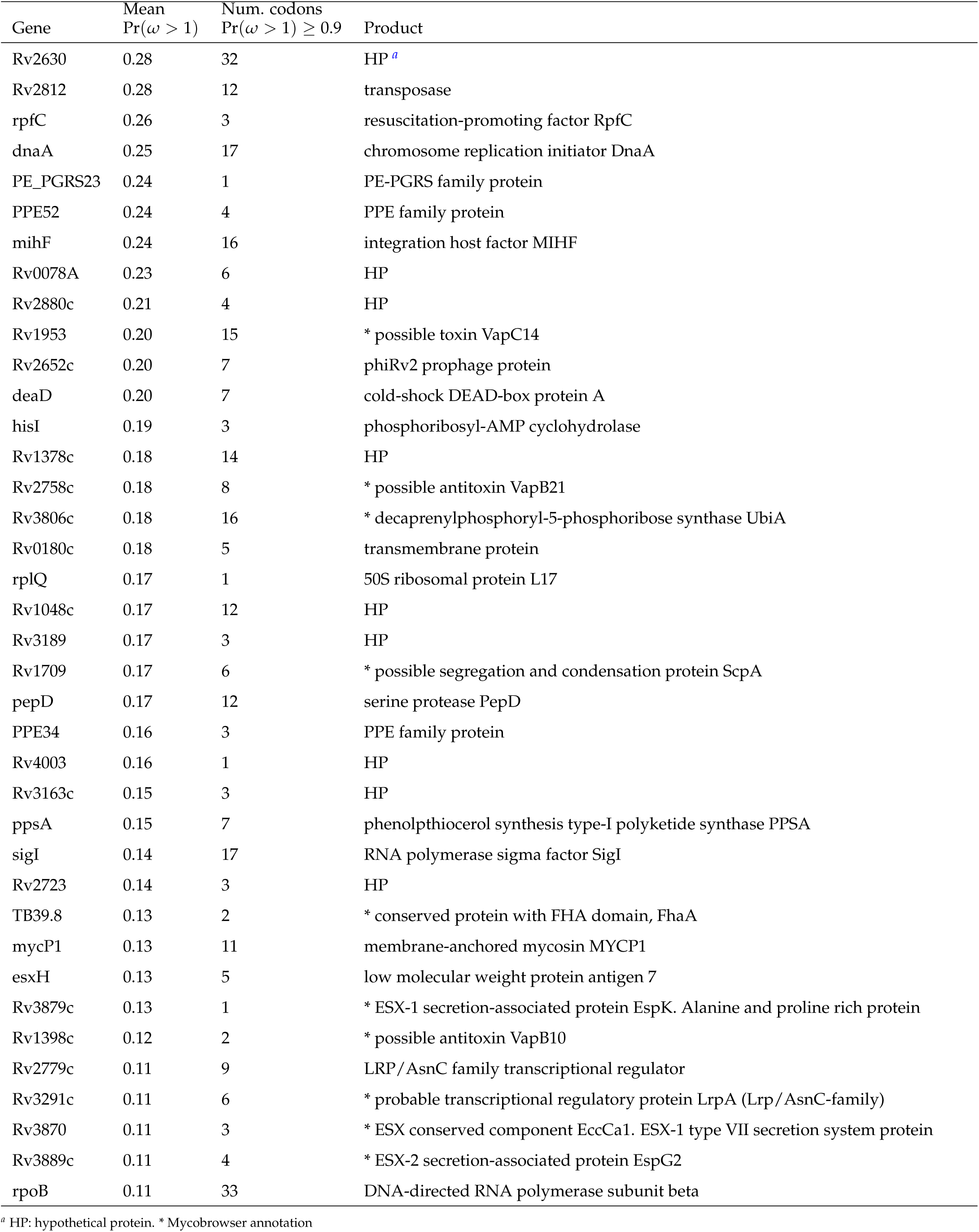
*Mycobacterium tuberculosis* genes with intermediate mean Pr(*ω >* 1) and individual codons with Pr(*ω >* 1) *≥* 0.9

| Gene | Mean<br>$\Pr(\omega > 1)$ | Num. codons<br>$\Pr(\omega > 1) \geq 0.9$ | Product |
| --- | --- | --- | --- |
| Rv2630 | 0.28 | 32 | HP <sup>a</sup> |
| Rv2812 | 0.28 | 12 | transposase |
| rpfC | 0.26 | 3 | resuscitation-promoting factor RpfC |
| dnaA | 0.25 | 17 | chromosome replication initiator DnaA |
| PE_PGRS23 | 0.24 | 1 | PE-PGRS family protein |
| PPE52 | 0.24 | 4 | PPE family protein |
| mihF | 0.24 | 16 | integration host factor MIHF |
| Rv0078A | 0.23 | 6 | HP |
| Rv2880c | 0.21 | 4 | HP |
| Rv1953 | 0.20 | 15 | * possible toxin VapC14 |
| Rv2652c | 0.20 | 7 | phiRv2 prophage protein |
| deaD | 0.20 | 7 | cold-shock DEAD-box protein A |
| hisI | 0.19 | 3 | phosphoribosyl-AMP cyclohydrolase |
| Rv1378c | 0.18 | 14 | HP |
| Rv2758c | 0.18 | 8 | * possible antitoxin VapB21 |
| Rv3806c | 0.18 | 16 | * decaprenylphosphoryl-5-phosphoribose synthase UbiA |
| Rv0180c | 0.18 | 5 | transmembrane protein |
| rplQ | 0.17 | 1 | 50S ribosomal protein L17 |
| Rv1048c | 0.17 | 12 | HP |
| Rv3189 | 0.17 | 3 | HP |
| Rv1709 | 0.17 | 6 | * possible segregation and condensation protein ScpA |
| pepD | 0.17 | 12 | serine protease PepD |
| PPE34 | 0.16 | 3 | PPE family protein |
| Rv4003 | 0.16 | 1 | HP |
| Rv3163c | 0.15 | 3 | HP |
| ppsA | 0.15 | 7 | phenolphthiocerol synthesis type-I polyketide synthase PPSA |
| sigI | 0.14 | 17 | RNA polymerase sigma factor SigI |
| Rv2723 | 0.14 | 3 | HP |
| TB39.8 | 0.13 | 2 | * conserved protein with FHA domain, FhaA |
| mycP1 | 0.13 | 11 | membrane-anchored mycosin MYCP1 |
| esxH | 0.13 | 5 | low molecular weight protein antigen 7 |
| Rv3879c | 0.13 | 1 | * ESX-1 secretion-associated protein EspK. Alanine and proline rich protein |
| Rv1398c | 0.12 | 2 | * possible antitoxin VapB10 |
| Rv2779c | 0.11 | 9 | LRP/AsnC family transcriptional regulator |
| Rv3291c | 0.11 | 6 | * probable transcriptional regulatory protein LrpA (Lrp/AsnC-family) |
| Rv3870 | 0.11 | 3 | * ESX conserved component EccCa1. ESX-1 type VII secretion system protein |
| Rv3889c | 0.11 | 4 | * ESX-2 secretion-associated protein EspG2 |
| rpoB | 0.11 | 33 | DNA-directed RNA polymerase subunit beta |

<sup>a</sup> HP: hypothetical protein. \* Mycobrowser annotation

**Table 3.** *Mycobacterium tuberculosis* genes with low mean Pr(*ω >* 1) and individual codons with Pr(*ω >* 1) *≥* 0.9

| Gene | Mean<br>$\Pr(\omega > 1)$ | Num. codons<br>$\Pr(\omega > 1) \geq 0.9$ | Product |
| --- | --- | --- | --- |
| Rv0585c | 0.11 | 8 | * probable conserved integral membrane protein |
| lprM | 0.11 | 2 | MCE-family lipoprotein LprM |
| serB1 | 0.11 | 13 | phosphoserine phosphatase |
| frdB | 0.10 | 2 | fumarate reductase iron-sulfur subunit FrdB |
| Rv0263c | 0.10 | 14 | HP <sup>a</sup> |
| sugI | 0.10 | 23 | sugar-transport integral membrane protein SugI |
| rpoC | 0.10 | 17 | DNA-directed RNA polymerase subunit beta' |
| parA | 0.09 | 10 | chromosome partitioning protein ParA |
| Rv2280 | 0.09 | 2 | dehydrogenase |
| Rv3885c | 0.09 | 3 | * ESX conserved component EccE2. ESX-2 type VII secretion system protein |
| Rv1277 | 0.09 | 1 | HP |
| cysD | 0.08 | 11 | sulfate adenylyltransferase subunit 2 |
| cstA | 0.08 | 5 | carbon starvation protein A CstA |
| rpfB | 0.08 | 6 | resuscitation-promoting factor rpfB |
| murD | 0.07 | 13 | UDP-N-acetylmuramoyl-L-alanyl-D-glutamate synthetase |
| Rv0634c | 0.07 | 5 | * possible glyoxalase II (hydroxyacylglutathione hydrolase) (GLX II) |
| Rv0842 | 0.07 | 3 | * probable conserved integral membrane protein |
| Rv0104 | 0.07 | 8 | HP |
| Rv2315c | 0.07 | 3 | HP |
| Rv0373c | 0.07 | 6 | carbon monoxide dehydrogenase large subunit |
| cyp130 | 0.06 | 5 | cytochrome P450 130 CYP130 |
| rpoA | 0.06 | 5 | DNA-directed RNA polymerase subunit alpha |
| Rv2824c | 0.06 | 6 | HP |
| Rv1273c | 0.06 | 5 | drugs-transport transmembrane ATP-binding protein ABC transporter |
| lysA | 0.05 | 2 | diaminopimelate decarboxylase LysA |
| Rv1130 | 0.05 | 25 | * possible methylcitrate dehydratase PrpD |
| pth | 0.05 | 2 | peptidyl-tRNA hydrolase |
| folC | 0.05 | 4 | bifunctional folylpolyglutamate synthase/dihydrofolate synthase FolC |
| ileS | 0.05 | 14 | isoleucyl-tRNA synthetase |
| gyrB | 0.04 | 4 | DNA gyrase subunit B |
| glcB | 0.04 | 5 | malate synthase G |
| Rv1795 | 0.04 | 8 | * ESX conserved component EccD5. ESX-5 type VII secretion system protein |
| Rv3679 | 0.04 | 1 | anion transporter ATPase |
| embB | 0.03 | 1 | indolylacetylinoitol arabinosyltransferase |
| Rv1702c | 0.03 | 10 | HP |
| ppp | 0.03 | 1 | serine/threonine phosphatase |
| Rv3802c | 0.02 | 5 | * probable conserved membrane protein |
| gyrA | 0.02 | 7 | DNA gyrase subunit A |
| Rv3428c | 0.01 | 2 | transposase |

<sup>a</sup> HP: hypothetical protein. \* Mycobrowser annotation

### Positive Selection in Known Resistance-Determining Genes

Figure 4 shows in detail the variation in *ω* along ten genes, ordered by the mean Pr(*ω >* 1) (and cross-referenced above Figure 3). The signature of selection in *rpoB*, which encodes RNA polymerase subunit *β*, exemplifies the evolutionary response to antibiotic usage. Subunit *β* is targeted by the first-line drug rifampicin, which binds the RNA polymerase, interfering with transcription of DNA to mRNA (see e.g. Palomino and Martin 2014). Strong evidence of positive selection is found in a 28-codon hotspot covering codons 427–454 coinciding with the *rifampicin resistance determining region* and including the common serine-to-leucine substitution at position 450 (S450L; positions relative to NC_000962.2). The population harbors a large number of alternative amino acid alleles in this region, represented by an accumulation of orange points in Figure 4; this provides the signature of elevated *d*_*N*_/*d*_*S*_. The extremely large sample size greatly enhances the ability to discover these alternative alleles, many of which are rare. For example, codon 445, which showed the highest point estimate of *ω* = 37.2, harbors 14 alleles encoding 12 different amino acids, with H445Y the most abundant amino acid substitution at only 1.5% frequency. Additional signals were observed in three peaks covering codons 44–45, 399–400 and 491. None of these sites is included in the WHO-endorsed GeneXpert MTB/RIF assay despite evidence of involvement in MDR-TB outbreaks (e.g. Makhado *et al.* 2018).

**Figure 4.**
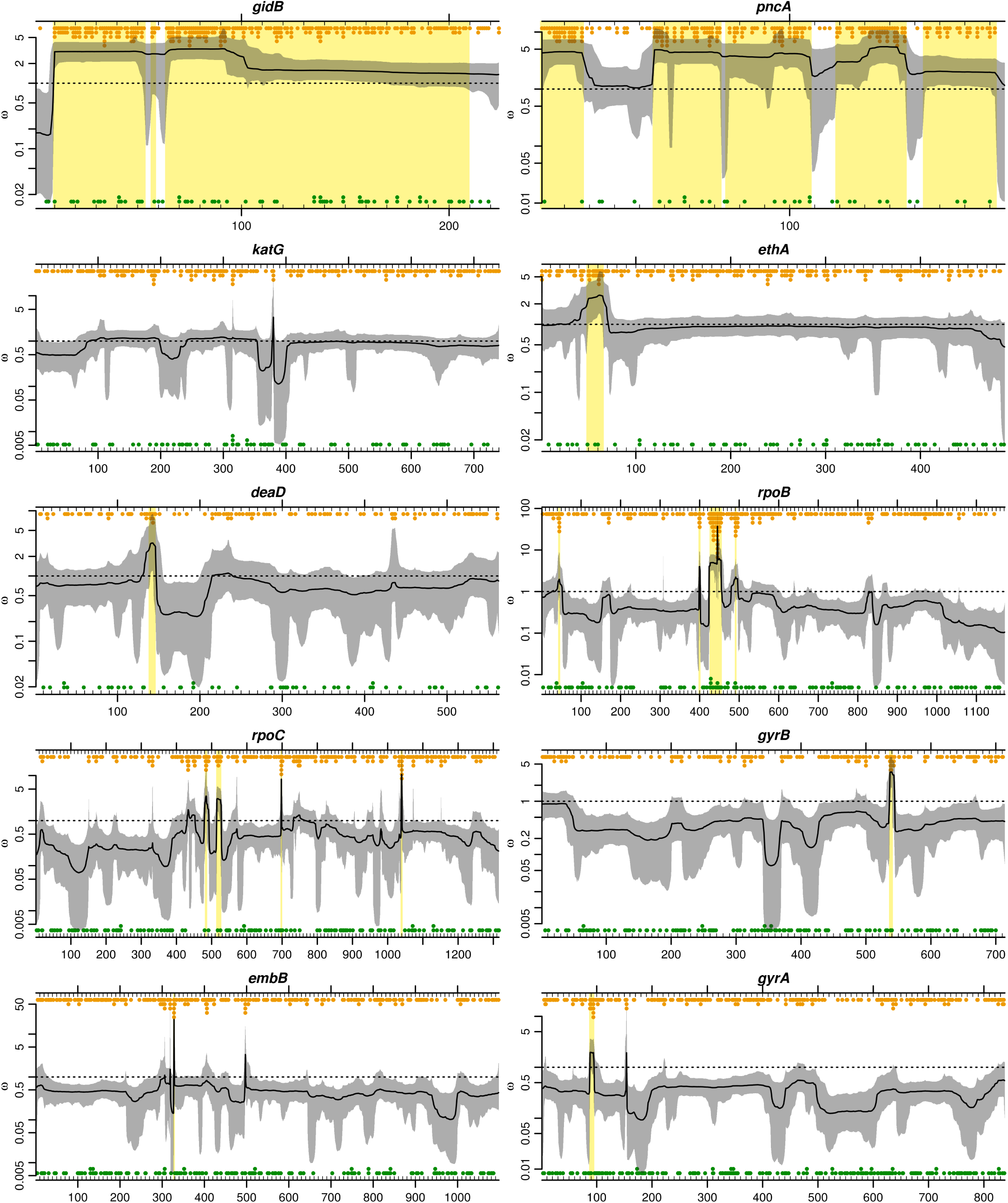
Evidence of positive selection in ten *Mycobacterium tuberculosis* genes across 10,209 genomes. Genes are ordered by the mean Pr(*ω >* 1) across codons, from highest (*gidB*) to lowest (*gyrA*). Point estimates (solid lines) and 95% credibility intervals (grey regions) for *ω* are shown across codons. Codons for which Pr(*ω >* 1) ≥ 0.9 are highlighted with yellow boxes. Stacked points indicate the number of alleles that are non-synonymous (orange) or synonymous (green) relative to the commonest allele.

The adjacent *rpoC* gene, encoding RNA polymerase subunit *β′*, showed similar peaks of positive selection against a backdrop of strong constraint. The regions showing strong evidence of positive selection covered codons 483–485, 515–525, 698 and 1039– 1040. Two of these regions coincide with high-probability compensatory mutations identified by Comas *et al.* (2012): V483A/G, D485H/N and N698H/K/S. The compensatory mutations mitigate the fitness deficit imposed on rifampicin-resistant *M. tuberculosis* by mutations in the rifampicin resistance determining region of *rpoB*. All three positions localize to the interface between RNA polymerase subunits *α* and *β′*, suggesting they play a role in the interaction between subunits (Comas *et al.* 2012). The extremely large sample size revealed other rare amino acid alleles at these positions that could also be compensatory: D485Y and N698D/L.

The World Health Organization (2018) report that 82% of rifampicin-resistant tuberculosis cases are also resistant to the first-line drug isoniazid, making them multidrug resistant tuberculosis (MDR-TB), which requires longer treatment with more toxic drugs. Isoniazid is a prodrug requiring activation by catalase-peroxidase, encoded by *katG*. Despite detecting the highest level of homoplasy in *katG* among 23 resistance-associated genes in a previous study of 2,099 genomes (Walker *et al.* 2015), *genomegaMap* did not detect evidence of positive selection surpassing the posterior probability theshold of 90%. The resistance-conferring S315T substitution, which Walker *et al.* (2015) found emerged 180 times, had a 78.5% probability of positive selection. However, in contrast to *rpoB* S450L in the rifampicin resistance determining region, *katG* S315T is surrounded by conserved sites. In the sliding window model used by *genomegaMap*, this dilutes the signal of S315T and weakens the evidence for positive selection. *GenomegaMap* also ignores the signal of homoplasy because it does not use a phylogenetic tree. In the case of *katG* S315T, these properties can be regarded as weaknesses of the approach, despite their advantages in other respects.

Resistance to the first-line drug ethambutol is conferred by mutations in *embB*, which encodes an essential part of the cell wall biosynthetic pathway (Palomino and Martin 2014). Selection is predominantly conservative in *embB*, with a single codon found to exhibit strong evidence of position selection, D328F/G/H/I/F. Position M306I/L/V, which has been implicated in ethambutol resistance, had a posterior probability of positive selection of only 56.1%, despite amino acid polymorphism. In fact, only two other codons in the entire gene, Q497H/K/P/R and Y319C/D/S, showed any evidence of positive selection (81.9% and 64.5% respectively). This demonstrates the strong constraint pervasive in *embB* and underlines the difficulty of detecting positive selection at sites whose neighbors are strongly conserved.

The DNA gyrase-encoding genes *gyrA* and *gyrB* display strong signatures of positive selection localized to the quinolone resistance determining regions, surrounded by strong constraint characteristic of essential proteins. A single region in each gene reached the 90% probability threshold, covering codons 88–94 in *gyrA* and 537–540 in *gyrB*. Several of these positions are known to confer resistance to second-line quinolone drugs, including *gyrA* A90E/G/V and D94A/G/H/N/Y (Palomino and Martin 2014).

Selection at *ethA*, which encodes a non-essential monooxygenase, appears dominated by neutral evolution, reminiscent of the general signature in *katG* whose product is also nonessential. Loss-of-function mutations in *ethA* prevent activation by monooxygenase of the second-line ethionamide from a prodrug to its active form (Palomino and Martin 2014). Strong evidence for positive selection is apparent in *ethA*, localized to codons 49-65. Like *katG*, this suggests that although resistance-conferring loss-of-function mutations can occur throughout the gene, they tend not to. The apparent neutrality of much of *ethA* and *katG* may therefore be misleading, and might instead reflect a balance between antimicrobial-imposed positive selection for loss-of-function mutations conflicting with functional constraint favoring conservation of the gene products.

Rapidly evolving genes dominated by positive selection are rare in *M. tuberculosis*, and exemplified by *pncA*. This gene encodes the non-essential enzyme pyrazinamidase, which converts the first-line prodrug pyrazinamide to its active form. Resistance to PZA is achieved by loss-of-function mutations in *pncA* (Palomino and Martin 2014). Function-ablating missense and nonsense mutations have arisen very rapidly in response to the widespread use of pyrazinamide, and unlike *katG* and *ethA*, positive selection appears to have won out over functional constraint throughout most of the gene. The five regions where evidence for positive selection is weaker may be under stronger functional constraint in environments where expression of the gene is favored.

The *gidB* gene shows strong evidence of positive selection throughout almost its entire length. This gene encodes a methyl-transferase that increases resistance to the second-line drug streptomycin. Streptomycin inhibits protein synthesis by binding to the 16S rRNA component of the 30S ribosomal subunit, increasing mistranslation. Loss-of-function of the *gidB* methyltransferase is thought to alter methylation of a highly conserved 16S rRNA residue, preventing binding by streptomycin (Okamoto *et al.* 2007; Wong *et al.* 2011). Like in *pncA*, this mechanism creates a selection pressure favoring missense and nonsense mutations throughout the gene. However, the modest increase in resistance conferred by this mechanism and the current status of streptomycin as a relatively less-frequently used, second-line drug with strong side effects suggests there may be other selection pressures driving *gidB* loss-of-function.

### Positive Selection in a Cold-Shock Protein

I scanned the *genomegaMap* results for evidence of positive selection at genes in which the selective forces driving adaptation are unknown or incompletely understood. In particular, I looked for genes with the characteristic signature of positive selection against a backdrop of functional constraint. The *deaD* gene, encoding cold-shock DEAD-box protein A and also known as *csdA*, is one such example (Figure 4).

DEAD-box proteins are a large family of ATP-dependent RNA helicase proteins found in prokaryotes and eukaryotes that separate double-stranded RNA molecules in an energy-dependent manner. They are named after their highly conserved Asp-Glu-Ala-Asp (D-E-A-D) motif. DEAD-box proteins are involved in ribosome biogenesis, translation initiation and RNA decay, fundamental processes that must dynamically respond to changes in environment and stress (Linder and Fuller-Pace 2013).

In *Escherichia coli*, the DeaD/CsdA protein has been characterized as essential for ribosome formation during cold shock because it separates stable secondary RNA structures which form at low temperature (*Jones et al.* 1996). DeaD/CsdA is important for biogenesis of both the 30S and 50S ribosome subunits, conferring tolerance towards mutants of other regulators and ribosomal proteins (Moll *et al.* 2002; Charollais *et al.* 2004). DeaD/CsdA has also been found to control gene expression at temperatures relevant to the mammalian host, and for modulating the carbon storage regulatory (Csr) system, which globally regulates mRNA translation and turnover (Vakulskas *et al.* 2014). Strong evidence of positive selection in *M. tuberculosis deaD* was evident at codons 139–145 encoding the sequence TPGRMID in most of the genomes. This sequence corresponds to motif Ib, consensus sequence TPGRXXD, one of a series of highly conserved motifs that characterize DEAD-box proteins. Motif Ib overlaps a nine-residue alpha helix (*α*7) beginning at codon 140 in *M. tuberculosis*. Sengoku *et al.* (2006) characterized the structure of the *Drosophila melanogaster* DEAD-box protein Vasa in detail. They found that two RecA-like domains in the DEADbox protein core bind a single RNA strand and sharply bend it. The bend avoids a clash between the RNA and a ‘wedge’ formed by *α*7 when the RNA is single stranded, whereas the unbound strand of an RNA duplex would be predicted to clash with the *α*7 wedge, resulting in disrupted base-pairing.

The residues homologous to four codons in motif Ib directly interact with the bound RNA (Sengoku *et al.* 2006). These positions exhibited a single alternative amino acid allele each across the 10,209 genomes: T139P, G141D, R142P and D145H. Two of the remaining positions exhibited multiple alternative amino acid alleles – P140L/S and M143I/R/V – while I144 was invariant. No synonymous variation was seen across the motif. Despite the relatively abundant amino acid variation in the motif in terms of allele numbers, the frequency of all substitutions except M143I/R/V was extremely low, below 0.5%. The sensitivity of the *d*_*N*_/*d*_*S*_ ratio to allele numbers, irrespective of allele frequencies, was observed earlier in *rpoB*. The diversity of rare alleles could mirror the mode of selection in the *rpoB* rifampicin resistance determining region, in which any of a large collection of amino acid substitutions improve fitness in the presence of the drug.

The DEAD-box motif itself, covering codons 163–166 and responsible for RNA binding, ATP binding and interdomain interactions, was situated in a region of very strong conservation, with a mean probability of positive selection of 0.7%. This, together with the general conservation throughout the gene, suggests that the effect of substitutions in motif Ib might not be to knock out the function of DeaD, but to modify it in some way. For instance, by altering conformation in such a way as to change interactions with other molecules.

Given the functional characterization of DeaD, candidate drivers of adaptation in motif Ib may in some way inhibit ribosome biogenesis or translation by interfering with ribosomal proteins, rRNAs or amino acids through mutation, for example with reactive oxygen radicals produced by the immune response, conformational change, for example binding by an antibiotic, or changes in molecular availability, for example caused by nutrient deprivation, cold shock or other stress. In the case of drug resistance, the detection of localized positive selection against a backdrop of strong constraint in *deaD* provides valuable context for future GWAS searching for genetic variants responsible for the growing problem of drug resistant infections.

### Conclusions

The main advantages of *genomegaMap* for estimating *d*_*N*_/*d*_*S*_ ratios within species are (i) it is fast no matter how large the sample size and (ii) it accounts for recombination. These advantages were achieved by extending the Wilson *et al.* (2011) approximation to the distribution of allele frequencies under parent-dependent mutation models, and assuming independence between codons. Simulations showed good performance despite these approximations.

Among the benefits of the approach, haplotype information is not required and missing data is easily handled, making *genomegaMap* suitable for short-read exome data in diploids and haploids. The *genomegaMap* approach is to treat *d*_*N*_/*d*_*S*_ as a substitution parameter. In this light, it can be seen as a general, likelihood-based method for estimating substitution parameters within species under parent-dependent mutation models.

The approach has several limitations. Sites are assumed independent between codons but linked within codons. Despite this, simulations showed good performance when recombination was high and low. Thus it was possible to analyse 10,209 genomes from *M. tuberculosis*, an almost perfectly clonal organism. One disadvantage of the independence assumption is ignoring homoplasy. In the *katG* example, this led to the surprising result that despite high homoplasy, no site achieved Pr(*ω >* 1) ≥ 0.9. The effects of violating other assumptions including constant population size, no population structure and random sampling were not investigated. The importance of sampling cannot be overstated, with signatures of selection entirely dependent on the selection pressures experienced by the populations analysed. Perhaps the greatest limitation of *genomegaMap* is its use of the *d*_*N*_/*d*_*S*_ ratio to characterize natural selection. Within species, *d*_*N*_/*d*_*S*_ is expected to vary even in a constant environment, with ratios closer to one expected for younger variants not yet exposed to selection for so long (McDonald and Kreitman 1991). Further, the form of positive selection that best predicts a high *d*_*N*_/*d*_*S*_ ratio is diversifying selection, in which any amino acid is favored over the incumbent. Diversifying selection may be relatively limited, to arms races e.g. between host and pathogen, or to heterogeneous environments e.g. immunologically diverse hosts. The evolution of resistance to antibiotics since their introduction in the 1940s may resemble such a Red Queen scenario, particularly as exposure is likely to vary from host-to-host.

Examples from *rpoB* and *deaD* showed that the signal of elevated *d*_*N*_/*d*_*S*_ stems mainly from the abundance of alternative amino acid alleles, relative to the number expected under neutrality, and not from allele frequencies. Some of these alternative alleles were detected at frequencies below 0.5%, demonstrating the value of extremely large sample sizes. The sliding window model employed by *genomegaMap* gained power to detect selection when positively selected sites were clustered as in *rpoB* and *deaD*, but missed the key isoniazid resistance-conferring S315T substitution of *katG* which is surrounded by highly conserved sites. Despite these limitations, the relatively simple interpretation of *d*_*N*_/*d*_*S*_ ratios means the approach continues to hold a strong appeal. For such applications, *genomegaMap* helps accelerate the exploitation of big data for gaining new insights into evolution within species.

## Acknowledgments

I would like to thank Nicola De Maio for insightful comments. D.J.W. is a Sir Henry Dale Fellow, jointly funded by the Wellcome Trust and the Royal Society (grant no. 101237/Z/13/Z) and is a Big Data Institute Robertson Fellow. The CRyPTIC Consortium is supported by grants from the Bill and Melinda Gates Foundation (OPP1133541) and a Wellcome Trust/Newton Fund-MRC Collaborative Award (200205/Z/15/Z). F.A.D. was supported by the Imperial Biomedical Research Centre.

## Appendix A: Members of The CRyPTIC Consortium

Derrick W Crook, Timothy EA Peto, A Sarah Walker, Sarah J Hoosdally, Ana L Gibertoni Cruz, Joshua Carter, Clara Grazian, Sarah G Earle, Samaneh Kouchaki, Yang Yang, Timothy M Walker, Philip W Fowler and David A Clifton, University of Oxford; Zamin Iqbal and Martin Hunt, European Bioinformatics Institute; E Grace Smith, Priti Rathod, Lisa Jarrett and Daniela Matias, Public Health England, Birmingham; Daniela M Cirillo, Emanuele Borroni, Simone Battaglia, Arash Ghodousi, Andrea Spitaleri and Andrea Cabibbe, Emerging Bacterial Pathogens Unit, IRCCS San Raffaele Scientific Institute, Milan; Sabira Tahseen, National Tuberculosis Control Program Pakistan, Islamabad; Kayzad Nilgiriwala and Sanchi Shah, The Foundation for Medical Research, Mumbai; Camilla Rodrigues, Priti Kambli, Utkarsha Surve and Rukhsar Khot, P.D. Hinduja National Hospital and Medical Research Centre, Mumbai; Stefan Niemann, Thomas Kohl and Matthias Merker, Research Center Borstel; Harald Hoffmann, Nikolay Molodtsov and Sara Plesnik, Institute of Microbiology & Laboratory Medicine, IML red, Gauting; Nazir Ismail, Shaheed Vally Omar, Lavania Joseph and Elliott Marubini, National Institute for Communicable Diseases, Johannesburg; Guy Thwaites, Thuong Nguyen Thuy Thuong, Nhung Hoang Ngoc and Vijay Srinivasan, Oxford University Clinical Research Unit, Ho Chi Minh City; David Moore, Jorge Coronel and Walter Solano, London School of Hygiene and Tropical Medicine and Universidad Peruana Cayetano Heredá, Lima; George F Gao, Guangxue He, Yanlin Zhao, Aijing Ma and Chunfa Liu, China CDC, Beijing; Baoli Zhu, Institute of Microbiology, CAS, Beijing; Ian Laurenson and Pauline Claxton, Scottish Mycobacteria Reference Laboratory, Edinburgh; Anastasia Koch, Robert Wilkinson, University of Cape Town; Ajit Lalvani, Imperial College London; James Posey, CDC Atlanta; Jennifer Gardy, University of British Columbia; JimWerngren, Public Health Agency of Sweden; Nicholas Paton, National University of Singapore; Ruwen Jou, Mei-Hua Wu, Wan-Hsuan Lin, CDC Taiwan; Lucilaine Ferrazoli, Rosaline Siqueira de Oliveira, Institute Adolfo Lutz, Sao Paolo. Authors contributing to the CRyPTIC consortium are (in alphabetical order): Irena Arandjelovic (Institute of Microbiology and Immunology, Faculty of Medicine, University of Belgrade, Belgrade, Serbia), Angkana Chaiprasert (Faculty of Medicine Siriraj Hospital, Mahidol University, Thailand), Iñaki Comas (Instituto de Biomedicina de Valencia (IBV-CSIC). Calle Jaime Roig, Valencia, Spain; FISABIO Public Health, Valencia, Spain; CIBER in Epidemiology and Public Health, Madrid, Spain), Francis A Drobniewski (Imperial College, London, UK), Maha R Farhat (Harvard Medical School, Boston, USA), Qian Gao (Shanghai Medical College, Fudan University, Shanghai, China), Rick Ong Twee Hee (Saw Swee Hock School of Public Health, National University of Singapore, Singapore), Vitali Sintchenko (Centre for Infectious Diseases and Microbiology - Public Health, University of Sydney, Sydney, Australia), Philip Supply (Genoscreen, Lille, France; Univ. Lille, CNRS, Inserm, CHU Lille, Institut Pasteur de Lille, U1019 - UMR 8204 - CIIL - Centre d’Infection et d’Immunitá de Lille, F-59000 Lille, France) and Dick van Soolingen (National Institute for Public Health and the Environment (RIVM), Bilthoven, The Netherlands). 10 D.

